# A Chemical Biology Toolbox for the Study of Protein Methyltransferases and Epigenetic Signaling

**DOI:** 10.1101/260638

**Authors:** Sebastian Scheer, Suzanne Ackloo, Tiago S. Medina, Matthieu Schapira, Fengling Li, Jennifer A. Ward, Andrew M. Lewis, Jeffrey P. Northrop, Paul L. Richardson, H. Ümit Kaniskan, Yudao Shen, Jing Liu, David Smil, Minkui Luo, Jian Jin, Dalia Barsyte-Lovejoy, Kilian V. M. Huber, Daniel D. De Carvalho, Masoud Vedadi, Colby Zaph, Peter J. Brown, Cheryl H. Arrowsmith

## Abstract

Protein methyltransferases (PMTs) comprise a major class of epigenetic regulatory enzymes with therapeutic relevance. Here we present a collection of chemical probes and associated reagents and data to elucidate the function of human and murine PMTs in cellular studies. Our collection provides inhibitors and antagonists that together modulate most of the key regulatory methylation marks on histones H3 and H4, providing an important resource for modulating cellular epigenomes. We describe a comprehensive and comparative characterization of the probe collection with respect to their potency, selectivity, and mode of inhibition. We demonstrate the utility of this collection in CD4^+^ T cell differentiation assays revealing the remarkable potential of individual probes to alter multiple T cell subpopulations with important implications for T cell-mediated processes such as inflammation and immuno-oncology. In particular, we demonstrate a role for DOT1L in limiting Th1 cell differentiation and maintaining lineage integrity.

## INTRODUCTION

Epigenetic regulation of gene expression is a dynamic and reversible process that establishes and maintains normal cellular phenotypes, but contributes to disease when dysregulated. The epigenetic state of a cell evolves in an ordered manner during cellular differentiation and epigenetic changes mediate cellular plasticity that enables reprogramming. At the molecular level, epigenetic regulation involves hierarchical covalent modification of DNA and the histone proteins that package DNA. The primary heritable modifications of histones include lysine acetylation, lysine mono-, di- or trimethylation, and arginine methylation. Collectively these modifications establish chromatin states that determine the degree to which specific genomic loci are transcriptionally active (Mendenhall et al., 2013).

Proteins that “read, write and erase” histone (and non-histone) covalent modifications have emerged as druggable classes of enzymes and protein-protein interaction domains (Arrowsmith et al., 2012). Histone deacetylase (HDAC) inhibitors and DNA hypomethylating agents have been approved for clinical use in cancer and more recently, clinical trials have been initiated for antagonists of the BET bromodomain proteins (which bind to acetyllysine on histones), the protein methyltransferases EZH2, DOT1L and PRMT5, and the lysine demethylase LSD1 (Shortt et al., 2017). The development of this new class of epigenetic drugs has been facilitated by the use of chemical probes to link inhibition of specific epigenetic protein targets with phenotypic changes in a wide variety of disease models, thereby supporting therapeutic hypotheses (Arrowsmith et al., 2015).

Methylation of lysine and arginine residues in histone proteins is a central epigenetic mechanism to regulate chromatin states and control gene expression programs (Ribich et al., 2017;

Wesche et al., 2017; Wu et al., 2017). Mono-, di- or tri-methylation of lysine side chains in histones can be associated with either transcriptional activation or repression depending on the specific lysine residue modified and the degree of methylation. Arginine side chain methylation states include monomethylation and symmetric or asymmetric dimethylation (**Figure 1**). In humans two main protein families carry out these post-translational modifications of histones. The structurally related PR and SET domain containing enzymes (protein lysine methyltransferases (PKMT)) methylate lysine residues on histone ‘tails’, and the dimeric Rossman fold protein arginine methyltransferase (PRMT) enzymes modify arginine. DOT1L has the Rossman fold, but is a monomer and modifies a lysine on the surface of the core histone octamer within a nucleosome (as opposed to the disordered histone tail residues). Many of these proteins also methylate non-histone proteins, and even less is known about non-histone methylation signaling (Buuh et al., 2017; Hamamoto et al., 2015).

**Figure 1.**
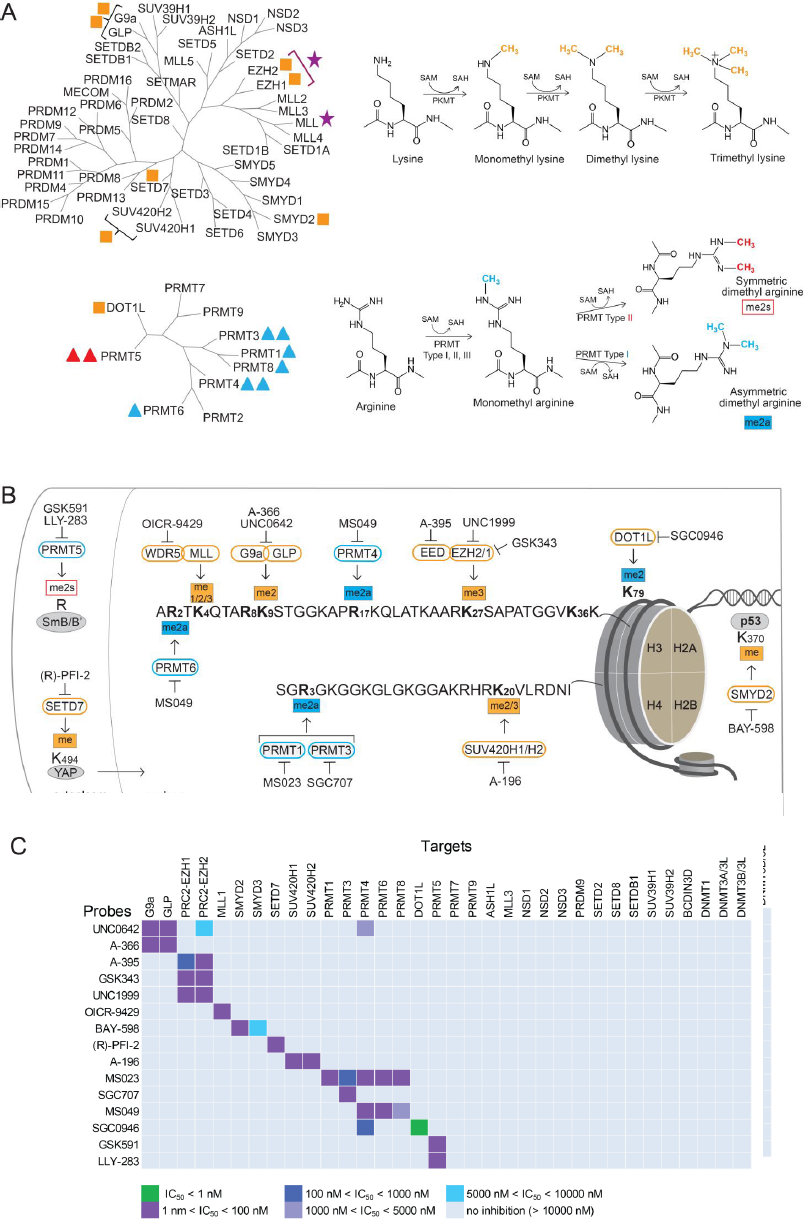
Summary of Chemical Probes. (A) Phylogenetic trees of human PR and SET domain lysine methyltransferases (upper tree), and the β-barrel fold enzymes (lower tree). Trees are annotated to show the chemical probes in this collection that inhibit PKMTs (orange square), monomethyl and asymmetric dimethyl PRMTs (blue triangle), symmetric dimethyl PRMTs (red triangle); and methyltransferase protein complexes (pink star). The number of annotations adjacent to each target is equal to the number of chemical probes for that target. See also **Figure 1B**. SAM = S-adenosylmethionine; SAH = S-adenosylhomocysteine (B) Detailed coverage of the major histone H3 and H4 methyl marks provided by this collection of chemical probes. The methylated lysine (K) and arginine (R) residues are annotated in bold font. The PMTs that write the marks are shown with orange (PKMTs) or blue (PRMTs) borders, along with the chemical probes that inhibit these PMTs. Also included are non-histone substrates (grey ovals) of PRMT5, SETD7, and SMYD2. See also **Figure 1A**. ‘me’ – methyl; ‘me2a’ – asymmetric dimethyl; ‘me2s’ – symmetric dimethyl; ‘me2/3’ – di- and trimethyl marks are written by the same enzyme. (C) Selectivity of each chemical probe has been assessed against 34 SAM-dependent methyltransferases. See also **Tables S1-S3**.

Here we describe a resource of chemical probes and related chemical biology reagents and knowledge to be used to probe the cellular function of PMTs, and link inhibition of select PMTs to biological mechanisms and therapeutic potential. We summarize the key features of this set of reagents and demonstrate their utility as a collection to uncover previously unknown links between epigenetic regulators and T cell biology in both humans and mice.

## RESULTS

### A Suite of Chemical Probes with Broad Coverage of the Major Histone Methyl Marks

**Table 1** lists chemical probes for human protein methyltransferases (PMTs) and key characteristics of their activity. This collection provides significant coverage of the human histone lysine and arginine methyltransferase phylogenetic trees (**Figure 1A**), but more importantly includes modulators of the major regulatory histone methylation marks (**Figure 1B**). Key among these are H3K9me2, H3K27me3, H3K79me2, and H4K20me2/3, each of which are “written” exclusively by G9a/GLP, PRC2 complex (via EZH1/2), DOT1L and SUV420H1/2 enzymes, respectively. As such, the respective chemical probes for these enzymes are able to reduce the global levels of their resultant mark in cells as measured by western blot, ‘in-cell’ western, or immunofluorescence assays (Bromberg et al., 2017; Konze et al., 2013; Liu et al., 2013; Yu et al., 2012). Other histone marks such as H3K4me1/2/3 are written by multiple enzymes. Thus, a chemical probe such as OICR-9429, which disrupts the MLL1 complex, is more likely to have specific effects only at loci targeted by MLL1, and not necessarily all methylated H3K4 loci (Grebien et al., 2015). PMTs also have many non-histone targets (Buuh et al., 2017; Hamamoto et al., 2015), including transcription factors such as p53 (Deimling et al., 2017; Eggert et al., 2016; Ferry et al., 2017), estrogen receptor (Zhang et al., 2016; Zhang et al., 2013), and cytosolic signaling factors such as MAP3K2 (Fu et al., 2016). In addition to histone modification, arginine methylation plays an important role in the function of RNA-binding proteins, ribosome biogenesis and splicing (Gao et al., 2017; Ren et al., 2010; Wall and Lewis, 2017). Thus, this collection of chemical probes constitutes a broad resource to link enzyme activity to a wide range of epigenetic and non-epigenetic methylation-mediated signaling pathways and biology.

**Table 1:** Summary of chemical probes and their chemotype-matched controls for protein methyltransferases.

| Target | Probe | *in vitro* IC<sub>50</sub> or K<sub>d</sub> (nM) | Cell line: assay | IC<sub>50</sub> (nM) | Control |
| --- | --- | --- | --- | --- | --- |
| WDR5* | OICR-9429<br>(Grebien et al., 2015) | 64 | HEK293: disrupt interaction of WDR5<sup>§</sup> with MLL1 & RbBP5 | 223 & 458 | OICR-0547 |
| EED* | A-395<br>(He et al., 2017) | 34 | RD: ↓ H3K27me3 | 90 | A-395N |
| EZH2 | GSK343<br>(McCabe et al., 2012) | 4 | HCC1806: ↓ H3K27me3 | 250 |  |
| EZH2,H1 | UNC1999<br>(Konze et al., 2013) | 10,45 | MCF10A: ↓ H3K27me3 | 124 | UNC2400 |
| DOT1L | SGC0946<br>(Yu et al., 2012) | 0.3 | MCF10A,A431: ↓ H3K79me2 | 10,3 | SGC0649 |
| G9a & GLP | A-366<br>(Sweis et al., 2014) | 4 & 38 | PC3: ↓ H3K9me2 | 300 | Use UNC0642 |
|  | UNC0642<br>(Liu et al., 2013) | <3 | PC3: ↓ H3K9me2 | 130 | Use A-366 |
| SUV420H1/2 | A-196<br>(Bromberg et al., 2017) | 21 | U2OS: ↓ H4K20me2/3 | 262/370<br>(SUV420H1) | A-197,<br>SGC2043 |
| SMYD2 | BAY-598<br>(Eggert et al., 2016) | 27 | HEK293, SMYD2<sup>§</sup>:<br>↓ p53K370me | 58 | BAY-369 |
| SETD7 | (R)-PFI-2<br>(Barsyte-Lovejoy et al., 2014) | 2 | MEFs and MCF7: ↑ nuclear YAP | sub-μM | (S)-PFI-2 |
| Type 1 PRMTs | MS023<br>(Eram et al., 2016) | 30,8<br>(PRMT1,6) | MCF7 (PRMT1), HEK293 (PRMT6<sup>§</sup>): ↓ H4R3me2a,<br>↓ H3R2me2a | 9,56<br>(PRMT1,6) | MS094 |
| PRMT3 | SGC707<br>(Kaniskan et al., 2015) | 31 | HEK293, PRMT3<sup>§</sup>:<br>↓ H4R3me2a | 91 | XY1 |
| PRMT4 | TP-064<sup>1</sup> | <10 | HEK293: ↓ Med12me2a,<br>↓ Baf155me2a | 43,<br>340 | TP-064N |
|  | †SKI-73<sup>2</sup> | 11 | MCF-7: ↓ Med12me2a | 540 | SKI-73N |
| PRMT4,6 | MS049<br>(Shen et al., 2016) | 44,63 | HEK293:<br>↓ Med12me2a (PRMT4),<br>↓ H3R2me2a (PRMT6<sup>§</sup>) | 1400,<br>970 | MS049N |
| PRMT5 | GSK591<br>(Duncan et al., 2016) | 11 | Z138: ↓ SmD3me2s | 56 | SGC2096 |
|  | LLY-283<sup>3</sup> | 22 | MCF7: ↓ SmBB'Rme2s | 30 | LLY-284 |

\*Subunits required for activity of their respective enzyme complexes.
§Cellular assay utilizes exogenous, transfected target
†SKI-73 is a pro-drug that is converted to the active compound by reductases in the cell.
↓ = decrease; ↑ = increase
<sup>1</sup>http://www.thesgc.org/chemical-probes/TP-064
<sup>2</sup>http://www.thesgc.org/chemical-probes/SKI-73
<sup>3</sup>http://www.thesgc.org/chemical-probes/LLY-283

### Chemical Probes are Potent, Active in Cells, and Highly Characterized for Selectivity

The development of these chemical probes was guided by principles practiced in the pharmaceutical industry to test the link between a specific protein and a putative biological or phenotype cellular trait (Arrowsmith et al., 2015; Blagg and Workman, 2017; Bunnage et al., 2013; Frye, 2010). A useful chemical probe should be reasonably potent in cells, selective for the intended target protein, and free of confounding off-target activities.

The chemical probes described here were each discovered using a biochemical enzymatic assay for the respective recombinant protein, or in some cases the relevant recombinant multiprotein enzyme complex, where the probe has been demonstrated to have an on-target potency with IC_50_ < 100 nM (**Table 1**). Each probe was evaluated in a customized cellular assay that tested the ability of the probe to reduce the level of methylation on its substrate in cells. All probes have significant, on-target cellular activity at 1 μM making them useful tools for cellular studies. Importantly these chemical probes are highly selective for their target protein (**Figure 1C**); each has been screened against a collection of up to 34 human SAM-dependent histone, DNA and RNA methyltransferases. The chemical probes within the lysine methyltransferase family are highly selective with measurable cross reactivity only seen between very closely related proteins such as G9a and GLP, or SUV420H1 and SUV420H2. Selectivity within the PRMT family, however, is more difficult to achieve, and a greater degree of cross reactivity is seen in this subfamily. The probes had minimal or no off-target activities when screened against a panel of 119 membrane receptors and ion channels and kinases (**Table S1**).

Importantly, most chemical probes are accompanied by a structurally similar control compound that has similar physicochemical properties but is inactive or much less active on its target enzyme (**Tables 1 and S2**). These inactive compounds are to be used alongside the active probe to control for unanticipated off-target activity of their common chemical scaffold. In cases where an appropriate control molecule could not be identified, an alternative strategy is to use multiple chemical probes with different chemotypes that inhibit the same target (**Tables 1 and S2**). Many of the PMTs discussed here have two or more chemical probes with different chemotypes, mechanisms of action, and/or potency profile. By using multiple diverse chemical probes that inhibit the same target, the user can build confidence in the link between cellular phenotype and inhibition of a specific target.

### Multiple Mechanisms of Inhibition Offer a Rich Source of Selective Chemical Probes

The methyl-donating cofactor S-adenosylmethionine (SAM) and methyl-accepting substrate of protein methyltransferases bind at juxtaposed but distinct sites that can each be targeted by small molecule inhibitors (**Figure 2**). Early efforts to inhibit PMTs focused on targeting the SAM binding site in analogy to targeting the ATP binding site of kinases (Copeland et al., 2009). However, it has been challenging to identify cell-penetrant compounds that bind to the polar SAM binding pocket and no universal methyltransferase inhibitor scaffold or “warhead” has yet been identified.

**Figure 2.**
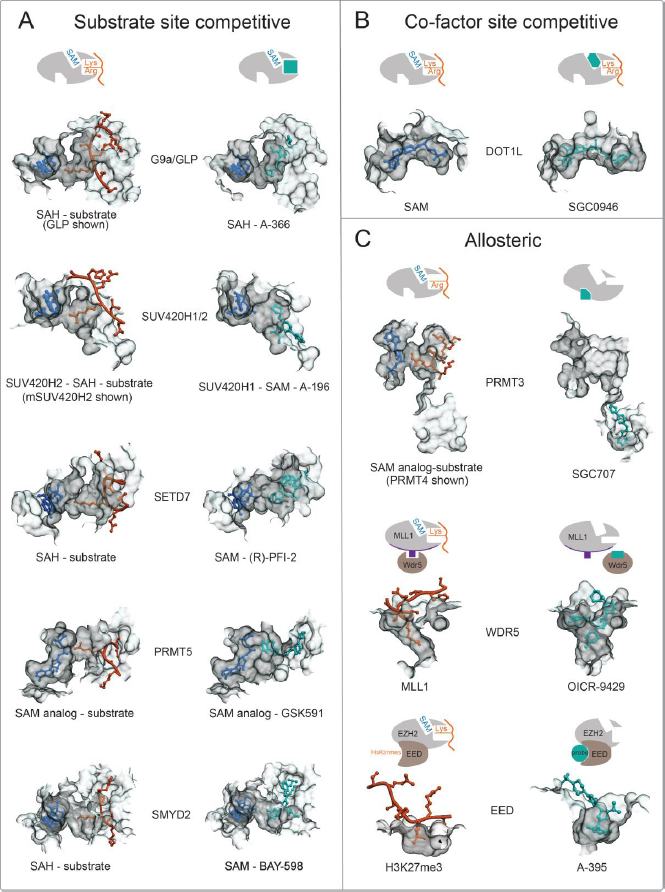
Structural Mechanisms of PRMT Inhibition by Chemical Probes. (A) Inhibitors of G9a (PDB: 3HNA (GLP with H3K9 substrate) and 4NVQ (A-366)); SUV420H2 (PDB: 4AU7 ( mSUV420H2 with H4 substrate) and SUV420H1 (PDB: 5CPR (A-196)); SETD7 (PDB: 1O9S (H3 substrate) and 4JLG (PFI-2)); PRMT5 (PDB: 4GQB (H4 substrate) and 5C9Z (GSK591)); SMYD2 (PDB: 3TG5 (p53 substrate) and 5ARG (BAY-598)) all bind in the substrate (peptide) binding pocket. (B) SGC0946 binds in the SAM-binding pocket of DOT1L thereby preventing cofactor binding (PDB ID: 3QOW (SAM), 4ER6 (SGC0946)). (C) Three distinct modes of allosteric inhibition of protein methyltransferases. SGC707 binds to PRMT3 in an allosteric site that prevents productive conformation of the enzyme’s activation helix (PDB ID: 4RYL). OICR-9429 binds to WDR5 and inhibits MLL1 activity by disrupting WDR5-MLL1-RBBP5 complex (PDB ID: 4QL1). A-395 binds to the EED subunit of the PRC2 complex thereby preventing binding of activating peptides (PDB ID: 5K0M).

The most frequent mode of inhibition for PMTs is binding of the probe within the substrate pocket, thereby preventing substrate binding (**Figure 2A**) (Barsyte-Lovejoy et al., 2014; Bromberg et al., 2017; Duncan et al., 2016; Eggert et al., 2016; Sweis et al., 2014). The high selectivity profile of the SET-domain chemical probes is likely related to the high degree of substrate selectivity of these enzymes (Schapira, 2016). These substrate-competitive probes also take advantage of the structural malleability of the substrate-binding groove to remodel the substrate-binding loops for optimal fit (**Figure 2A**, grey contours). Interestingly, binding of many of these substrate competitive probes is also dependent on cofactor SAM, which in some cases directly interacts with the probe molecule, and also is known to help stabilize formation of the substrate-binding pocket of SET domain proteins (Schapira, 2016). These contributions of SAM can be a confounding factor when interpreting enzyme inhibitory kinetic data for these probes. In **Figure 2** we have focused on the mechanism of action of each probe based on the structures of their complexes with their target enzyme. The potent SAM competitive inhibitors, SGC0946 (Yu et al., 2012) (**Figure 2B**) and LLY-283, have adenosine-like moieties with hydrophobic substituents replacing the methionine of SAM, while UNC1999 (Konze et al., 2013) and GSK343 (Verma et al., 2012) have a novel pyridone-based scaffold that binds in and near the unique pocket of the PRC2 multiprotein complex (Brooun et al., 2016).

A third class of inhibitors has allosteric mechanisms that can induce long-range structural perturbations, or take advantage of sites of protein-protein interactions within multi-subunit PMT complexes. The allosteric inhibitor SGC707 (Kaniskan et al., 2015) occupies a pocket that is 15 Å away from the site of methyl transfer, but prevents formation of the catalytically competent conformation of the PRMT3 dimer (**Figure 2C**, top panel). OICR-9429 (Grebien et al., 2015) and A-395 (He et al., 2017) are antagonists that make use of the peptide binding pocket in the essential WD40 subunits (WDR5 and EED) of the multiprotein MLL1 and PRC2 complexes, respectively (**Figure 2C**, middle and bottom panels). OICR-9429 binds in the central pocket of WDR5 preventing the latter’s interactions with MLL and histone peptides, resulting in diminished methyltransferase activity of the complex (Grebien et al., 2015). A-395 binds in the analogous pocket of EED, thereby preventing its interaction with trimethylated peptides that activate the PRC2 holoenzyme (He et al., 2017). Both chemical probes burrow into the central cavity of their respective WD40 protein targets occupying a remodeled binding pocket that is significantly larger than that of the peptides which they replace. Thus, a common feature revealed by the co-crystal structures of these probes with their target protein is the significant remodeling of the substrate or allosteric binding sites to selectively accommodate each probe molecule.

A caveat of targeting scaffolding subunits of chromatin complexes is that such proteins are often components of multiple complexes, with diverse functions in cells. For instance, WDR5 interacts with at least 64 different proteins, including the oncoprotein c-MYC, and disrupting the WDR5 protein interaction network has diverse functional consequences beyond the MLL complex. In this regard, chemical probes are ideal tools to investigate the biochemical and cellular outcome of targeting a specific protein interaction interface. Taken together, this collection of chemical probes reveals the wide array of mechanisms by which PMTs may be inhibited.

### Chemical Probe Reagents to Study Cellular Specificity and Interaction Networks

Chemical proteomics represents a powerful approach to investigate the molecular target profile of chemical probes (Huber and Superti-Furga, 2016). In total, ten affinity reagents were synthesized for seven targets. Of these reagents, three have been previously verified for target selectivity (Barsyte-Lovejoy et al., 2014; Grebien et al., 2015; Konze et al., 2013), and for an additional five probes (for four targets) we indicate the recommended site of derivatization (**Table S2**). These suggestions are based on empirical data obtained during each probe discovery program, including target/probe co-crystal structures and chemical structure-activity-relationships (SAR). Using three of these new affinity reagents we generated cellular selectivity data for the DOT1L inhibitor SGC0946, EED antagonist A-395, and the Type I PRMT inhibitor MS023 (Eram et al., 2016). All three affinity probes were able to specifically enrich their cognate targets from cell lysates as demonstrated by immunoblotting. This enrichment could be competed with the respective underivitized “free” chemical probe suggesting a specific interaction not affected by immobilisation (**Figure S1**). Profiling of the pan Type I PRMT inhibitor MS023 in HEK293 cell lysates using label-free quantification (LFQ) protein mass spectrometry showed engagement of several PRMTs including PRMT1, 3, 4, and 6 and their respective binding partners (**Figure 3A,D**). PRMT8, which MS023 is known to inhibit, was not detected. This is consistent with lack of expression of the neuronal-specific PRMT8 in HEK cells. The negative control compound MS094 had no effect on PRMT target engagement or interactome. Similarly, investigation of the EED antagonist A-395 in G401 cell lysates revealed several well-known PRC2 complex members which were co-purified with the cognate target EED (**Figure 3B,E**) similar to results obtained with a structurally distinct EED antagonist (Qi et al., 2017). As expected, pre-treatment with the negative control A-395N even at high concentration did not affect binding of EED and the other PRC2 complex members to the A-395 affinity matrix, confirming its applicability as an additional tool for EED and PRC2. Chemoproteomic analysis of the DOT1L inhibitor SGC0946 in Jurkat cell lysates indicated that the probe is highly specific for DOT1L, confirming earlier results obtained with HL-60 cells (Gilan et al., 2016) (**Figure 3C**).

**Figure 3.**
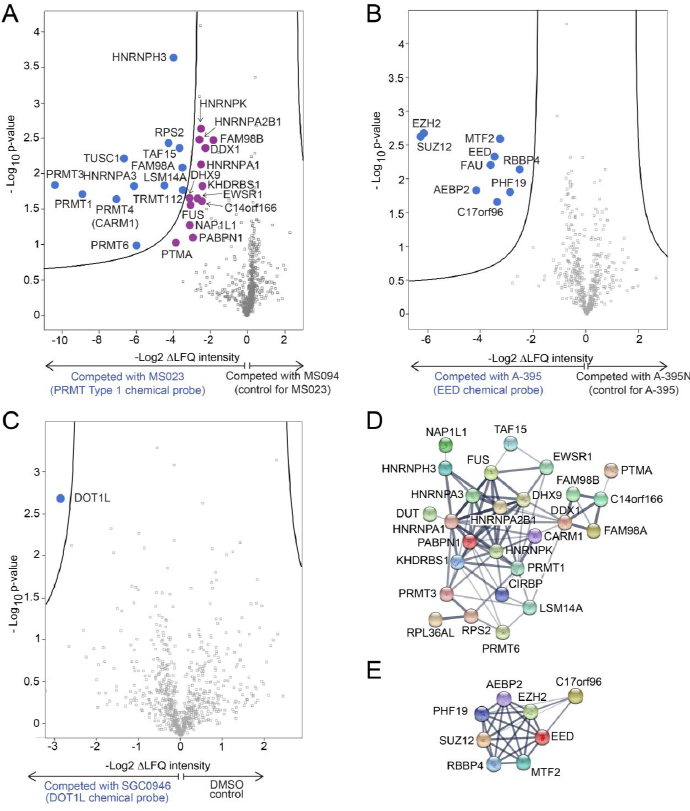
Affinity Reagents for Chemoproteomics. (A) Volcano plot of MTM7172-enriched proteome (labelled blue) from HEK293 cell lysate, with targets significantly (FDR=0.05, S0 = 0.2) competed by 20 µM MS023 or MS094, with respect to DMSO control. Proteins marked in purple indicate known interactors of the identified direct targets of MS023 close to the significance threshold. (B) Volcano plot of (A-395)-biotin enriched proteome (labelled blue) from G401 cell lysate, with targets significantly (FDR=0.05, S0 = 0.2) competed by 20 µM A-395 or A-395N, with respect to DMSO control. (C) Volcano plot of SGC2077-enriched proteome (labelled blue) from Jurkat cell lysate, with targets significantly (FDR=0.05, S0 = 0.2) competed by 20 µM SGC0946, with respect to DMSO control. (D) STRING network evaluation of targets significantly competed by MS023. Lines in the STRING evaluation represent evidenced interactions, with line thickness indicating confidence (high to low). (E) STRING network evaluation of targets significantly competed by A-395. See also **Figure S1**.

### Approaches to Use the Collection

Each chemical probe can be used to investigate the effects of inhibiting the respective PMT in a biochemical or cellular assay. Thus, each probe and its control(s) will be valuable tools in hypothesis-driven research projects. We envision, however, that these tools will be equally valuable in prospective, hypothesis-generating discovery screens to identify specific PMTs whose inhibition is linked with a certain phenotype. To date, much of the research using epigenetic inhibitors has focused on cancer research, with inhibitors of several PMT targets progressing into clinical trials. While these oncology studies are of great interest and medical importance, we sought to systematically investigate the landscape of PMT inhibition in a non-cancer setting, namely CD4^+^ T helper (Th) cell differentiation.

### Regulation of Th Cell Differentiation by PMTs

Cellular differentiation is guided by chromatin-mediated changes in gene expression in response to extracellular cues. In the immune system, naïve Th cells can adopt a wide range of cellular fates depending upon the external signals received during activation by antigen-presenting cells (O’Shea and Paul, 2010). For example, interleukin (IL)-12 promotes the development of Th1 cells that express the transcription factor T-bet (Tbx21) and produce interferon (IFN)-γ, while activation in the presence of IL-4 leads to Th2 cells that express GATA-3 and secrete IL-4, IL-5 and IL-13. In addition, Th17 (RORγt-expressing and IL-17A-producing) and induced regulatory T (Treg) cells (FOXP3-expressing) develop from naïve Th cells in the presence of TGFβ and IL-6 or TGFβ alone, respectively. The magnitude of the immune response to a given stimulus is dependent on the abundance and relative proportions of Th subtypes, and dysregulation of the balance among Th subtypes contributes to diseases such as autoimmunity and cancer (Zhu and Paul, 2008). Previous studies using gene-deficient mice have identified central roles for PMTs including MLL1 (Yamashita et al., 2006), G9a (Antignano et al., 2014; Lehnertz et al., 2010) and EZH2 (Tumes et al., 2013; Yang et al., 2015) in Th cell differentiation and stability.

We tested the effect of inhibiting each methyltransferase on Th cell differentiation, focusing first on IFN-γ-producing Th1 cells (**Figures 4A, B, S2, S3**). Isolated naïve CD4^+^ Th cells from the spleen and peripheral lymph nodes of mice with a fluorescent reporter of IFN-γ expression (Reinhardt et al., 2009) (IFN-γ-YFP mice) were activated for four days under Th1 polarizing conditions in the presence of each probe and its respective control compound (when available). Treatment with UNC1999 or GSK343 which target the catalytic subunits of PRC2 (EZH1/2), (but not the control compound UNC2400) significantly increased the expression and production of the signature cytokine IFN-γ under Th1 polarizing conditions. These results are consistent with previous studies showing that T cell-specific deletion of EZH2 resulted in enhanced Th1 cell differentiation in mice (Tumes et al., 2013; Yang et al., 2015; Zhang et al., 2014). We also observed a significant increase of IFN-γ producing CD4^+^ T cells using A-395 (but not A-395N), an antagonist of the EED subunit of PRC2 which prevents the latter’s enzymatic activity (He et al., 2017), demonstrating that the enhanced IFN-γ expression and production was independent of the chemotype of the active probe, further strengthening the link between PRC2 inhibition and Th1 cell differentiation. Our results also identified a role for the H3K79 methyltransferase DOT1L in regulation of Th1 cell differentiation. DOT1L inhibition resulted in an increase in the frequency of viable IFN-γ^+^ CD4^+^ cells and higher production of IFN-γ relative to the control compound and Th1 cells alone (**Figures 4A, S3A, B**). To further confirm that these observed differentiation phenotypes were related to inhibition of the methyltransferase activity of PRC2 and DOT1L we performed western blot analyses of the histone methyl marks deposited by each enzyme. Th1 cells treated with UNC1999 or A-395 (but not with the control compounds UNC2400 or A-395N, respectively) showed a significant reduction of H3K27me3 relative to untreated cells. Similarly, western blot analysis of H3K79me2 in the presence of SGC0946 showed an almost complete global loss of H3K79me2, but showed no change with the inactive SGC0649 or with the inhibitors of the PRC2 complex (A-395 or UNC1999) (**Figure 4E**). Thus, inhibition of specific histone methyltransferases and loss of specific histone methylation marks was associated with phenotypic changes in mouse Th cells.

**Figure 4.**
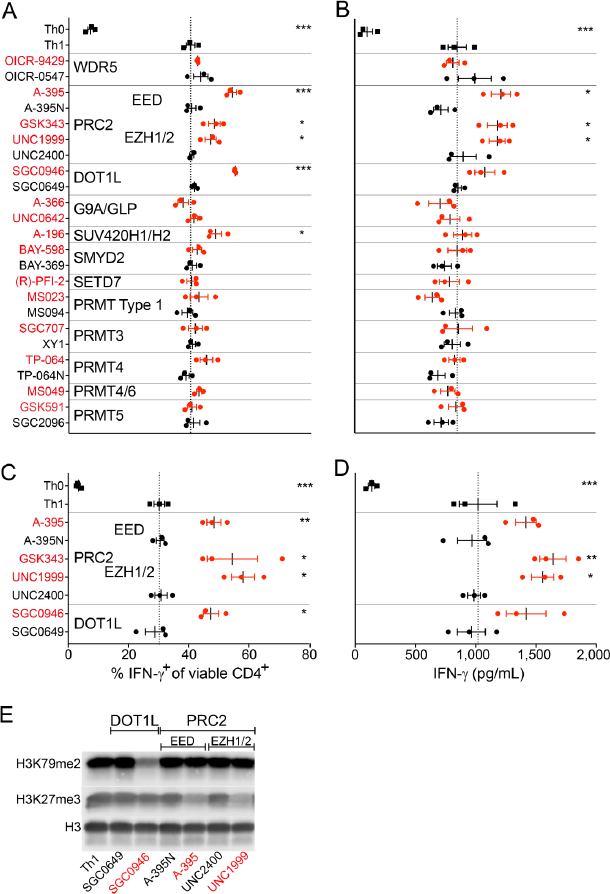
Differential Effects of PMT Inhibition on Murine and Human Th1 Cell Differentiation. (A, B) CD4^+^ T cells from the spleen and peripheral lymph nodes of IFN-γ-YFP reporter mice were enriched and polarized under Th1 cell conditions in the absence (Th0) or presence of indicated probes (1 μM; red) or their controls (where available; black). (A) FACS analysis after 4 days for intracellular YFP reporter signal (representing IFN-γ expression). (B) Secreted IFN- γ was analyzed by ELISA in the supernatant of the same experiment. Each data point represents one of three biological replicates and the data shown is representative of three independent experiments. (C, D) CD4^+^ T cells from the blood of three healthy human donors were cultured under Th1 cell-polarizing conditions in the presence or absence of indicated probes or their controls. (C) FACS analysis after 4 days for intracellular IFN-γ. (D) Secreted IFN-γ was analyzed by ELISA in the supernatant of the same experiment. Each data point represents one of three donors. Dotted lines visualize the mean frequency of IFN-γ-positive Th1 cells in the absence of probes (A, C) or the mean concentration of IFN-γ in the supernatant (B, D). Significant differences are indicated with an asterisk and were calculated using one-way ANOVA (*p ≤ 0.05, **p ≤ 0.01, ***p ≤ 0.001). (E) Western blot analysis of the effect of indicated inhibitors on the trimethylation of H3K27 and dimethylation of H3K79 in CD4^+^ T cells under Th1 polarizing conditions. Data shown is representative of 2 independent experiments. See also **Figures S2 and S4**.

### PMT-Mediated Regulation of Th1 Cell Responses is Conserved in Humans

To determine whether human T cells responded in a manner similar to mouse T cells, we activated human naïve peripheral blood CD4^+^ T cells for 4 days in the presence of Th1 cell-polarizing conditions. Consistent with our results for murine Th cells, inhibition of PRC2 (UNC1999, GSK343, A-395) and DOT1L (SGC0946) potentiated the effects of Th1 cell activation, resulting in a higher frequency of IFN-γ-producing Th1 cells as well as increased IFN-γ production, while control probes had no effect (**Figures 4C, D, S4**). These results identify a novel and central role for PRC2 and DOT1L in limiting Th1 cell differentiation in mice and humans.

### DOT1L-Dependent H3K79me2 is Dynamically Regulated During Th1 Cell Differentiation

Since to our knowledge DOT1L has not been examined in Th cell differentiation and function, we further investigated this enzyme using our mouse reporter system. In this four-day polarization assay, we only observed an enhancement of IFN-γ production with DOT1L inhibition if the cells were treated starting from day 0 or 1 of the culture, but not if the cells were exposed to the probe for only the last 1 or 2 days of polarization (**Figure S3C**). This is consistent with literature showing that reduction of histone methyl marks is dependent on the duration of exposure to methyltransferase inhibitors, often requiring days of exposure (Daigle et al., 2011; Vedadi et al., 2011). To explore the dynamics of H3K79me2 during Th1 cell differentiation, we monitored the reduction of H3K79me2 and the production of IFN-γ over a period of 4 days (**Figure 5**). Western blot analysis of global H3K79me2 shows that the mark is reduced in Th cells following activation in the presence of the control compound SGC0649 (**Figure 5A)**, suggesting that Th cell activation alone leads to a reduction of H3K79me2 independent of DOT1L inhibition. However, inhibition of DOT1L by addition of SGC0946 resulted in further reduction of H3K79me2 by day 2 that was maintained until day 4 post-activation (**Figure 5A**). This earlier and increased reduction of H3K79me2 induced by SGC0946 correlated with heightened IFN-γ levels (**Figure 5B**). These data establish a correlation between reduction of the H3K79me2 mark and enhanced production of IFN-γ and validate that reduced H3K79me2 correlates with increased production of IFN-γ under Th1 cell-polarizing conditions. To assess whether DOT1L inhibition affected T cell proliferation, we used flow cytometric tracking of naïve Th cells labelled with a fluorescent dye (CFSE), and stimulated under either neutral (Th0) or Th1 cell-polarizing conditions in the presence of SGC0946 or SGC0649. We observed no effect on T cell proliferation (**Figure S3D**), suggesting that DOT1L inhibition likely affects the Th1 cell differentiation program without altering proliferative capacity.

**Figure 5.**
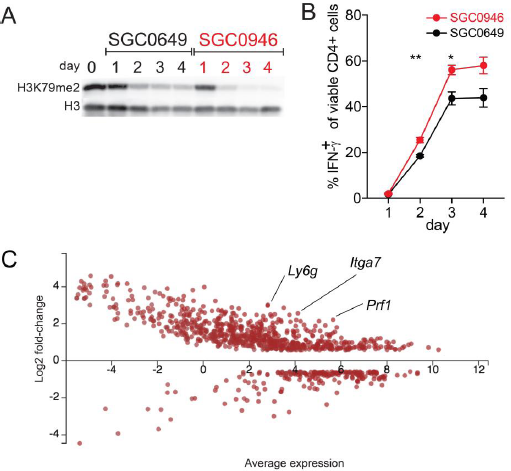
DOT1L-Dependent H3K79me2 is Dynamically Regulated at Lineage-Specific Promoters. A) Analysis of H3K79me2 in CD4+ T cells under Th1 polarizing conditions in the presence of the DOT1L inhibitor SGC0946 (1 µM) or the control compound SGC0649 (1 µM) with time as measured by western blot. (B) Analysis of IFN-γ production by FACS under the same conditions as in (A). Significant differences are indicated with an asterisk and were calculated using one-way ANOVA (*p ≤ 0.05, **p ≤ 0.01). Data shown is representative of three independent experiments. (C) DOT1L-dependent regulation of gene expression in Th1 cells. MA^#^-plot comparing SGC0649-treated and SGC0946-treated IFN-γ^+^ CD4^+^ T cells after Th1 polarization of up and down genes. Dots represent genes that show >1.5-fold increase (FDR 0.01) or decrease in SGC0946-treated CD4^+^ T cells compared to SGC0649-treated CD4^+^ T cells. Average expression is the mean of 3 biological replicates. See also **Figure S3**. #M (log ratio); A (mean average)

We next carried out unbiased gene expression analysis. Naïve Th cells from IFN-γ-YFP mice were stimulated under Th1 cell-polarizing conditions for 4 days in the presence of SGC0946 or SGC0649. IFN-γ^+^ CD4^+^ Th1 cells were purified by cell sorting, and RNA was isolated for RNA-Seq analysis. Comparing SGC0946- and SGC0649-treated IFN-γ-positive Th1 cells, we observed 750 genes that were significantly upregulated, with 208 genes downregulated when DOT1L was inhibited (**Figure 5C**). We observed that inhibition of DOT1L led to expression of non-canonical genes for Th1 cells including perforin 1 (*Prf1*), α7 integrin subunit (*Itga7*), and Ly6G (*Ly6g*). Thus, our data suggest that DOT1L-dependent mechanisms are potentially important for limiting Th1 cell differentiation and maintaining lineage integrity.

### PMTs Differentially Regulate Th Cell Differentiation

We next extended our analysis of the probe collection to identify PMTs that may be involved in differentiation of not only Th1, but also Th2, Th17 or Treg cells. We activated naïve Th cells from either wild-type C57BL/6 mice or transgenic reporter mice (IFN-γ-YFP, IL-4-GFP, or FOXP3-EGFP) under Th1, Th2, Th17 or Treg cell differentiation conditions for four days in the absence or presence of PMT probes or control compounds, and analysed expression of lineage-specific markers. As summarized in **Figure 6**, we found a wide range of effects on T cell differentiation both between different probes as well as between different stimulation conditions for a single probe or chemotype. For example, inhibition of DOT1L and the PRC2 complex (EZH1/2, EED) enhanced effector cell differentiation (Th1, Th2 and Th17) with no effect on Treg cell development, while inhibition of G9a/GLP (UNC0642, A-366), strongly affected Treg cell differentiation. These data are consistent with our previous results demonstrating a key role for G9a in promoting Treg cell function (Antignano et al., 2014; Lehnertz et al., 2010).

**Figure 6.**
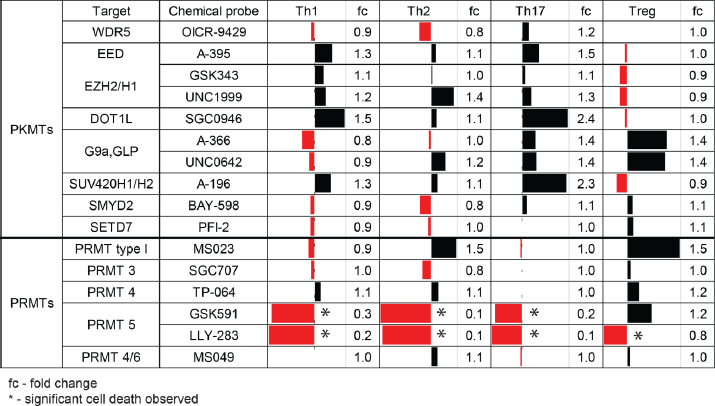
Overview of PMT Inhibition on Murine CD4+ T cell Differentiation. Each bar represents the mean percentage of up- (black) or down-regulation of the signature cytokines IFN-γ, IL-4 (lower panel) or IL-13 (upper panel), IL-17 or the transcription factor Foxp3 under Th1, Th2, Th17 and Treg cell-differentiation conditions, respectively. Data shown is from 2 (lower panel) or 3 (upper panel) individual experiments with 3 biological replicates each. Numbers show fold change of treatment with probe (1 μM) relative to the control (1 μM) or to DMSO (in the absence of control). See also **Figure S5**.

Another chemical probe that showed strong promotion of Treg cell differentiation is MS023 which inhibits the type I PRMT enzymes, PRMT1, 3, 4, 6, and 8. PRMT8 is a neuronal-specific enzyme and not expressed in T cells (Penney et al., 2017). MS023 was not tested on PRMT2 because the latter has not yet been demonstrated to be an active methyltransferase enzyme (Lakowski and Frankel, 2009). Comparison of this result with that of the more selective probes for PRMT3 (SGC707), PRMT4 (TP-064) and PRMT4/6 (MS049) all of which had little or no effect on Treg differentiation, suggests that the primary effect of MS023 is due to inhibition of PRMT1. This screen suggests new potential strategies for promotion of Treg cell differentiation through inhibition of type I PRMTs, or PRMT1, should a selective PRMT1 probe become available. Such compounds may be beneficial to subdue aberrant inflammatory immune responses.

Inhibition of PRMT5 using peptide competitive (GSK591) and SAM competitive (LLY-283) chemical probes clearly down-regulated the signature cytokines for effector cell differentiation, but also significantly increased cell death (**Figure S5**). PRMT5 has been shown to be required for growth and viability of rapidly proliferating cells such as cancer cell lines (Gulla et al., 2017). Although the mechanisms are not fully understood, PRMT5 knockdown slows the cell cycle in NIH3T3 cells and induces G1 arrest in 293T and MCF7 cells (Scoumanne et al., 2009; Wei et al., 2012). It is possible that similar mechanisms are at play in Th cells. The effect of PRMT5 on Treg cell differentiation is unclear since the chemical probes yielded opposite results (even across 6 biological replicates).

## DISCUSSION

Here we present a collection of chemical biology reagents to modulate cellular methylation signaling, especially chromatin-mediated signaling. Each probe has been characterized for its selectivity within the human SET and PRMT methyltransferase families. Most of our chemical probes are accompanied by structurally similar inactive compounds that serve as negative controls for potential off-target effects of their common chemical scaffold. We have annotated the chemical structure of each chemical probe showing where they can be chemically derivatized to create additional reagents such as biotinylated probes for chemo-proteomics. Using several examples of such derivatives we demonstrate the cellular selectivity for their targets. Importantly, these probes and related compounds may be used for research without restrictions.

Using our collection of PMT chemical probes, we identified several PMTs that differentially regulate Th cell differentiation, with both expected and new results. For example, inhibition of the PRC2 complex led to enhanced Th1, Th2 and Th17 cell responses and a reduction in Treg cell development, which is consistent with the phenotype observed in mice with a genetic deletion of EZH2 (Tumes et al., 2013). In addition, inhibition of G9a resulted in a significant increase in the frequency of FOXP3-expressing Treg cells, which is in agreement with our previous results (Antignano et al., 2014). Our unbiased screen also identified novel regulators of Th cell differentiation. Inhibition of DOT1L promoted effector Th cell responses with little effect on Treg cell differentiation, as did type I PRMT inhibition. While the potential enhancement of a regulatory immune response by inhibitors of G9a or by members of the Type I PRMT family may be beneficial to control immune responses in inflammatory disease, an increase of pro-inflammatory responses by inhibiting the PRC2 complex or DOT1L might be beneficial to boost the immune response in underperforming immune systems. Thus, the PMT chemical probe library provides a novel toolbox to examine the biological role of methylation-mediated signaling in cellular assays.

Among the targets in our chemical probe collection, several have agents currently in clinical trials for cancer (EZH2, EED, DOT1L, and PRMT5) and additional PMT inhibitors are in preclinical studies. The availability of well characterized, unencumbered chemical probes to such targets will enable the research community to better understand the mechanisms and consequences of PMT inhibition in a wide variety of cellular contexts providing important new knowledge to help guide clinical development, patient stratification, and novel applications of PMT inhibitors. Of particular interest are the pro-inflammatory activities of PRC2 and DOT1L inhibitors, which may be beneficial in the oncology setting.

Overall, our collection of PMT chemical probes, control compounds, and chemical biology reagents will be an outstanding resource to identify key epigenetic regulators of any cellular phenotype. Additional uses of the collection may include (i) synthetic lethal screens of the probes versus genetic ablation of individual genes, (ii) screens for optimal combinations of PMT probes with existing standard of care therapies in cellular disease models, and (iii) investigation of endogenous PMT protein complexes in a variety of cells without the need to introduce tagged exogenous protein ‘baits’.

## SIGNIFICANCE

1. A collection of chemical probes, their chemotype-matched controls, and related chemical biology reagents enable research in methylation-dependent epigenetic regulation.
2. Detailed descriptions of the potency, selectivity and structural mechanisms of inhibition of each chemical probe as well as demonstration of how to derivatize each probe for creation of new reagents.
3. Demonstration of the utility of the chemical probe collection to modulate T cell differentiation, revealing a novel role for DOT1L inhibitor SGC0946 in promoting Th1 cell differentiation.

## ACKNOWLEDGEMENTS

We thank our numerous medicinal chemistry collaborators in the pharmaceutical industry, the Ontario Institute for Cancer Research, and the Jin, Frye, and Luo Labs who helped develop probes in this resource. Michael Curtin and Hongyu Zhou (Abbvie) contributed to the development of biotinylated A-395. Kazuhide Nakayama (Takeda) contributed to the development of TP-064 and TP-064N. 7TM, kinase, and ion channel off-target selectivity screening was kindly supplied by Eurofins-Cerep. Additional Ki determinations and receptor binding profiles were generously provided by the National Institute of Mental Health’s Psychoactive Drug Screening Program, Contract # HHSN-271-2013-00017-C (NIMH PDSP). The NIMH PDSP is directed by B.L. Roth MD, PhD at the University of North Carolina at Chapel Hill and Project Officer J. Driscoll at NIMH, Bethesda, Maryland, USA. We would like to thank B. Kessler, S. Bonham and R. Rischer from the TDI Discovery Proteomics Facility for their support. This work was supported by the Canadian Institutes of Health Research [FDN-154328 and 128090 to CHA, and FDN-148430 and 201512MSH-360794-228629 to DDC], Canadian Cancer Society [CCSRI 703716] to DDC, the Australian National Health and Medical Research Council (project grants 1104433 and 1104466 to CZ), and the Structural Genomics Consortium (SGC) which is a registered charity (number 1097737) that receives funds from AbbVie, Bayer Pharma AG, Boehringer Ingelheim, Canada Foundation for Innovation, Eshelman Institute for Innovation, Genome Canada through Ontario Genomics Institute [OGI-055], Innovative Medicines Initiative (EU/EFPIA) [ULTRA-DD grant no. 115766], Janssen, Merck KGaA, Darmstadt, Germany, MSD, Novartis Pharma AG, Ontario Ministry of Research, Innovation and Science (MRIS), Pfizer, São Paulo Research Foundation-FAPESP, Takeda, and Wellcome. TM is supported by “Conselho Nacional de Desenvolvimento Cientȷfico e TecnolɃgico (CNPq)”, Brazil. CZ is a VESKI Innovation Fellow. JAW, AML and KVMH are grateful for funding by Myeloma UK. J.J. acknowledges the support by the grants R01GM122749, R01CA218600, and R01HD088626 from the U.S. National Institutes of Health.

